# Measuring mutagenicity in ecotoxicology: A case study of Cd exposure in *Chironomus riparius*

**DOI:** 10.1101/2020.11.02.365379

**Authors:** Halina Binde Doria, Ann-Marie Waldvogel, Markus Pfenninger

**Affiliations:** LOEWE Centre for Translational Biodiversity Genomics, Senckenberg Biodiversity and Climate Research Centre, Georg-Voigt-Str. 14-16, D-60325, Frankfurt am Main, Germany; Department of Molecular Ecology, Senckenberg Biodiversity and Climate Research Centre, Georg-Voigt-Str. 14-16, D-60325, Frankfurt am Main, Germany; Department of Ecological Genomics, Institute of Zoology, University of Cologne, Zülpicher Straße 47b, D-50674 Cologne, Germany; Institute for Molecular and Organismic Evolution, Johannes Gutenberg University, Johann-Joachim-Becher-Weg 7, D-55128, Mainz, Germany

**Keywords:** chronic exposure, non-biting midge, mutagenicity assay

## Abstract

Existing mutagenicity tests for metazoans lack the direct observation of enhanced germline mutation rates after exposure to anthropogenic substances, therefore being inefficient. Cadmium (Cd) is a metal described as a mutagen in mammalian cells and listed as a group 1 carcinogenic and mutagenic substance. But Cd mutagenesis mechanism is not yet clear. Therefore, in the present study, we propose a method coupling short-term mutation accumulation (MA) lines with subsequent whole genome sequencing (WGS) and a dedicated data analysis pipeline to investigate if chronic Cd exposure on *Chironomus riparius* can alter the rate at which *de novo* point mutations appear. Results show that Cd exposure did not affect the basal germline mutation rate nor the mutational spectrum in *C. riparius*, thereby arguing that exposed organisms might experience a range of other toxic effects before any mutagenic effect may occur. We show that it is possible to establish a practical and easily implemented pipeline to rapidly detect germ cell mutagens in a metazoan test organism. Furthermore, our data implicate that it is questionable to transfer mutagenicity assessments based on *in vitro* methods to complex metazoans.

**Main find of the work:** Cd chronic exposure under environmental realistic concentrations did not exert mutagenicity; It is questionable to transfer *in vitro* mutagenicity assessments to complex metazoans.

## 1. INTRODUCTION

During the last decades, the introduction of genomics into ecotoxicology grew in importance together with stress-induced mutagenesis and how it can alter ecosystems (Bianchi et al., 2011; de Raat et al., 1985). But the current mutagenicity tests lack the efficiency and reliability regarding their evaluation of the mutagenic potential in metazoans. Therefore, method improvements in ecotoxicological testing and risk assessment procedures are needed for an improved description, measurement and evaluation of adverse effects in organisms after exposure to possibly mutagenic toxicants.

Mutagens and stress-induced mutagenesis are different from other toxicants and toxic effects, because they promote heritable changes in the genome of an organism by enhancing the rate at which mutations appear in the germline. They are also conceptually different from genotoxic effects, i.e. somatic mutations, which affect only the exposed individual. Thus, the harmful influence of germ line mutagens does not end when exposure ceases (Gibbons and LeBaron, 2017; Ram and Hadany, 2014; Sakumi, 2019). Many mutations do not affect organismal fitness, but if they do, they are generally deleterious with few exceptions (Eyre-Walker and Keightley, 2007). An artificially increased mutation rate can therefore lead to deterioration of population fitness, in particular in small populations and although rarely accelerating adaptive evolution, it has the potential to irreversibly change the evolutionary trajectory (de Raat et al., 1985; Galhardo et al., 2007; Lynch and Gabriel, 1990). But if mutation rates are truly affected by environmental and anthropogenic factors is a lingering, yet not fully understood, question (Bull et al., 2019).

The main reason for this knowledge gap is that the methods currently in use for this purpose have several limitations:

- First, *in vitro* approaches like the Ames test (Ames et al., 1973) rely on microorganisms. Significant differences in mutational and DNA repair mechanisms between unicellular and multicellular organisms offer different possibilities for substances and stress to interfere with replication fidelity (Morita et al., 2010; Sung et al., 2012). This is consistent with the observation that mutagenic effects of the same substance can vary among bacteria and eukaryotes (Bergkamp et al., 1988). Further, to increase test sensibility, the great majority of strains used present inactivated repair mechanisms (McCann et al., 1975). In combination with the different exposure situation of, e.g., unicellular microbes and invertebrates with complex tissue structure, the transferability of results is limited.
- Second, cell cultures do not take into account the complexity of multicellular organisms. (Fowler et al., 2012). This is especially true for the cell type most relevant for mutagenic changes, i.e. germ cells in metazoan organisms which depart early during organismal development, divide less often and have different mutation rates compared to somatic cells (Murphey et al., 2013; Walter et al., 1998).
- Third, current tests performed in multicellular complex organisms access rather genotoxicity than germline mutagenicity itself by analysing clastogenicity, aneuploidy and DNA fragmentation with the micronuclei or comet assays (Duez et al., 2003; Girardello et al., 2016).

Therefore, because genotoxicity and mutagenicity are not synonymous terms, a combined approach looking for *in vivo* genotoxicity of known *in vitro* mutagens, still lacks the direct observation of an enhanced germline mutation rate after exposure to putative mutagens. As a consequence, the current rating of substances with regard to their mutagenic potential in metazoan target organisms likely generated both false positive and negative assessments, *i.e.* substances were flagged as mutagenic that actually are not and *vice versa* (Benigni et al., 2010; Fowler et al., 2012; Zeiger, 2019). Both may have severe negative consequences for environmental protection, human welfare, legislation and the economy (Amiard-Triquet et al., 2015).

At present, no more than thirty chemical agents (only four of those are considered as environmental pollutants), have had their mutagenicity directly assessed with next generation sequencing (NGS) technology (Beal et al., 2019; Bull et al., 2019; Du et al., 2017; Wamucho et al., 2019). Each year, around 50,000 to 100,000 chemical substances are registered in the European Union. But, from the total of registered substances, around 100,000 different chemicals are already being used on the EU market. Nonetheless, only a small fraction of those has been evaluated regarding their health and environmental impacts (Goldenman et al., 2017). This highlights not only the need to more research in this field, but also the development of an effective and straightforward whole-organism metazoan mutagenicity assay in ecotoxicology. In the present study we describe the development of a recently introduced mutation rate test (Oppold and Pfenninger, 2017) to an effective ecotoxicological mutagenicity test tool for metazoan organisms. The test consists of a combination of short-term mutation accumulation (MA) lines, whole genome sequencing (WGS) and dedicated data analysis.

As proof of principle, we investigated the effects of chronic stress caused by Cadmium (Cd) exposure on the occurrence of *de novo* point mutations in *Chironomus riparius.* Cd is a non-essential highly toxic metal. It is considered to be a widespread pollutant in water bodies (Wang et al., 2019). Cd has long known clastogenic effects both *in vitro* and *in vivo* (Hartwig, 2010; Seoane and Dulout, 2001). Further, it was described as a mutagen in mammalian cells and has already been proven to inhibit DNA repair and promote oxidative stress, which might be linked to mutagenicity (Filipič et al., 2006; Sherrer et al., 2018). Therefore, Cd is listed as a group 1 carcinogenic and mutagenic substance (IARC, 2012). Yet, despite the evident genotoxicity, data on mutagenicity is conflicting and relies only on *in vitro* data (Hartwig, 2010). The non-biting midge *C. riparius* is distributed throughout Europe. It is used both by the US Environmental Protection Agency (EPA) the Organisation for Environmental Co-operation and Development (OECD) as a test species in ecotoxicological standard tests to be used for the environmental risk assessment of chemicals (OECD, 2010; USEPA, 1996). Its genome was recently sequenced and annotated, enabling more complex genomic studies (Oppold et al., 2017; Schmidt et al., 2020). Moreover, it is known that *C. riparius* can rapidly adapt to environmental stress (Nowak et al., 2009; Pfenninger and Foucault, 2020).

Our goal was to establish a practical pipeline which can be easily implemented and rapidly analysed to detect germ cell mutagens in metazoan test organisms and make recommendations about experimental design and sample allocation. In the long-term, this information can be useful for the preparation of an international standardized test guideline for mutagenicity.

## 2. MATERIAL AND METHODS

### 2.1. Test Compound and Chemical Analyses

Cadmium chloride salt (Merck, Germany) was diluted in deionized dechlorinate water and kept as stock solutions of 10 mg/L. Aqueous Cd concentrations at the end of the test were determined by inductively coupled plasma mass spectrometry (detection limit 0.0002 mg/L) (DIN EN ISO 17294-2; 2014). Cd concentrations on the sediment were assessed after extraction in a HCl + HNO3 mixture (aqua regia) (DIN EN 13657; 2003) and analysed by inductively coupled plasma optical emission spectrometry (detection limit of 0.2 mg/kg) (DIN EN ISO 11885; 2009).

### 2.2. Experimental Design

The present study adopted an approach already established and validated by Oppold and Pfenninger (2017) for the estimation of mutation rates that combines the advantages of MA lines with those of the trio approach. We adjusted the method to evaluate the mutagenic impact of Cd exposure over three generations in *C. riparius* (second to fifth generation).

A long-term laboratory strain of *C. riparius*, the “Laufer population”, from which the reference draft genome was sequenced (Oppold et al., 2017; Schmidt et al., 2020) was used. All MA lines originated from a single egg rope (F0) raised under optimal conditions to avoid selection. Subsequently to the successful reproduction of its offspring (F1), thirty egg-ropes were collected and used to establish the MA lines. After hatching, clutches were individually put in tall bowls (20 cm, 14,5 cm Ø) filled with a 1.5 cm sediment layer and 1.150 L of water where they completed their entire life cycle. Fifteen clutches were under control conditions and the remaining fifteen were exposed to 32 μg/L of Cd at the beginning of every generation. Only a single egg-clutch originating from each MA line was chosen to start the following next generation. Test vessels were kept at 22 ± 0.5 °C, 60% relative humidity, light: dark rhythm of 16:8 h photoperiod and were constantly aerated. Water evaporation in the test vessels was compensated by adding demineralized water. Conductivity was maintained between 550 - 650 μS/cm and pH around 8. All the lines were fed as described elsewhere (Foucault et al., 2019; Oppold and Pfenninger, 2017).

Due to swarm fertilization of females and the ensuing impossibility to determine the parents of a particular egg-clutch, the reference pool consisted of the collected adults from the first generation. At the end of the fifth generation, a single female individual from each of ten randomly chosen MA lines from each treatment was then collected and sent to sequencing.

### 2.3. Whole Genome Sequencing and Bioinformatic analysis

DNA was extracted with the Blood and Tissue QUIAGEN Kit according to manufacturer’s instructions. In order to obtain an ancestor to generate a baseline to identify *de novo mutations* (DNMs) 140 legs, one of each individual from F1, were pooled together to an expected mean coverage of 60X. After five generations one female of each of the MA-lines was whole-genome sequenced to an expected mean coverage of 30X on Illumina NovaSeq 6000 platform. Sequencing libraries were generated using NEBNext® DNA Library Prep Kit following manufacturer’s recommendations. The genomic DNA was randomly fragmented, then the DNA fragments were end polished, A-tailed, and ligated with the NEBNext adapter for Illumina sequencing, and further PCR enriched by P5 and indexed P7 oligos. The PCR products were purified (AMPure XP system) and resulted libraries were analyzed for size distribution by Agilent 2100 Bioanalyzer and quantified using real-time PCR. Reads were individually adapter clipped and quality trimmed, using Trimmomatic (Bolger et al., 2014).

The cleaned reads of the ancestor and the single female individuals from each MA line were processed according to the best practices of the GATK-pipeline (McKenna et al. 2010). Reads were mapped with bwa mem (v0.7.10–r789, Li and Durbin 2009) against the reference genome draft (NCBI: ERR2696325) (Schmidt et al., 2020) and filtered using a combination of Picard tools v1.123 (https://broadinstitute.github.io/picard/) to mark the duplicates and GATK v.3.3.0 (Mckenna et al., 2010) for realignment around indels and recalibration of bases. The resulting bam files obtained then an individual identifier of the sample (SM:) field in the read-group tag and were merged with Picard’s MergeSamFiles.

Unlike in Oppold & Pfenninger (2017), we used the software accuMUlate (Howell et al., 2018) to assess the Single Nucleotide Mutations (SNM). This tool has the advantage to perform basically identical analyses more or less automatically, however, at the cost of disregarding indel mutations. accuMUlate was then run using the reference genome and some individual parameters as input (Supplementary Material – S1).

The output table was then strictly filtered with a custom Python script according to the following parameters: probability of a mutation/one mutation/correct descendant genotype (>=0.90); number of reads matching the putatively-mutant allele in samples that are not mutants (=0); AD test statistic for mapping quality difference (>=1.95); p-value from a Fisher’s exact test for Stand Bias and Pair-mapping rate difference (>0.05). This step, as showed in Long et al. (2016) is decisive to filter out false positive and inconclusive mutations associated with mapping error and low coverage areas. Lastly, the resulting candidate list was then visually checked with IGV (Robinson et al., 2017) for final validation (Supplementary material – S2).

### 2.4. Data analysis - Bayesian framework

We used a Bayesian implementation of a Poisson test to check the difference in the rate of mutations per generational passage between control and Cd treatment (Billoir et al., 2008; Fox, 2010) (R library BayesianFirstAid,- Bååth, 2014). This implemented model uses an uninformative Jeffreys prior on lambda. The posterior distribution was sampled with default values (3 chains, 5000 iterations). In this context, the test assumes that the number of callable sites does not vary between control and treatment. We tested for this assumption with a Mann-Whitney U test.

We tested for differences in the mutational spectrum (transitions/transversions and AT> GC/GC>AT mutations) between control and treatment with a two-sided Chi-square test. The number of sites for which a mutation could be called and the μ estimation - ratio of mutations per number of callable sites over five generations per haploid genome - were calculated individually (Oppold and Pfenninger 2017).

## 3. Results

### 3.1. Experimental conditions and accuMUlate validation

Each generation took between 25 to 30 days. Cd concentrations in water at the end of the life-cycle was 1± 0.1528 μg/L. The Cd concentrations in the sediments was below the method’s detection limit (0.2 mg/kg). The raw data and mappings of all individuals and the parental pool can be found at ENA (study number: PRJEB40079). Mean sequence coverage ranged from 36.20x to 46.71x for the MA lines individuals and 75.61x for the reference pool. The number of callable sites ranged from 94,074,863 bp to 147,108,695 bp. There was no significant difference in the number of called sites between treatments and controls (Mann-Whitney U = 31, p = 0.45).

Two Cd MA lines were identified as mutator lines. Cd2 accumulated 24 SNMs, which was confirmed by resequencing a sibling. In MA line Cd9, more than 14,000 candidate mutations were called in F5 and 300 in the previous generation. Strong evidence suggests that no mismapping or sequencing error was involved in the high number of candidate mutation calls. Therefore, being likely true mutator lines, they were excluded from subsequent analysis.

Overall, we identified 37 mutations during 50 generational passages in the control 10 MA lines and 28 during 40 generational passages from 8 valid Cd treatment MA lines, with a range of 1 to 7 mutations per individual (Table 1). The respective rate estimates were 0.74 (95% HDI 0.52-1.0) mutations per generational passage for the control and 0.71 (0.46-0.98) for the Cd treatment (Figure 1). The rate ratio (control/treatment) was therefore 1.1. There was fairly equal evidence for the rate ratio being larger or smaller than 1 (42.1% < 1 < 52.7%, Figure 1).

**Figure 1.**
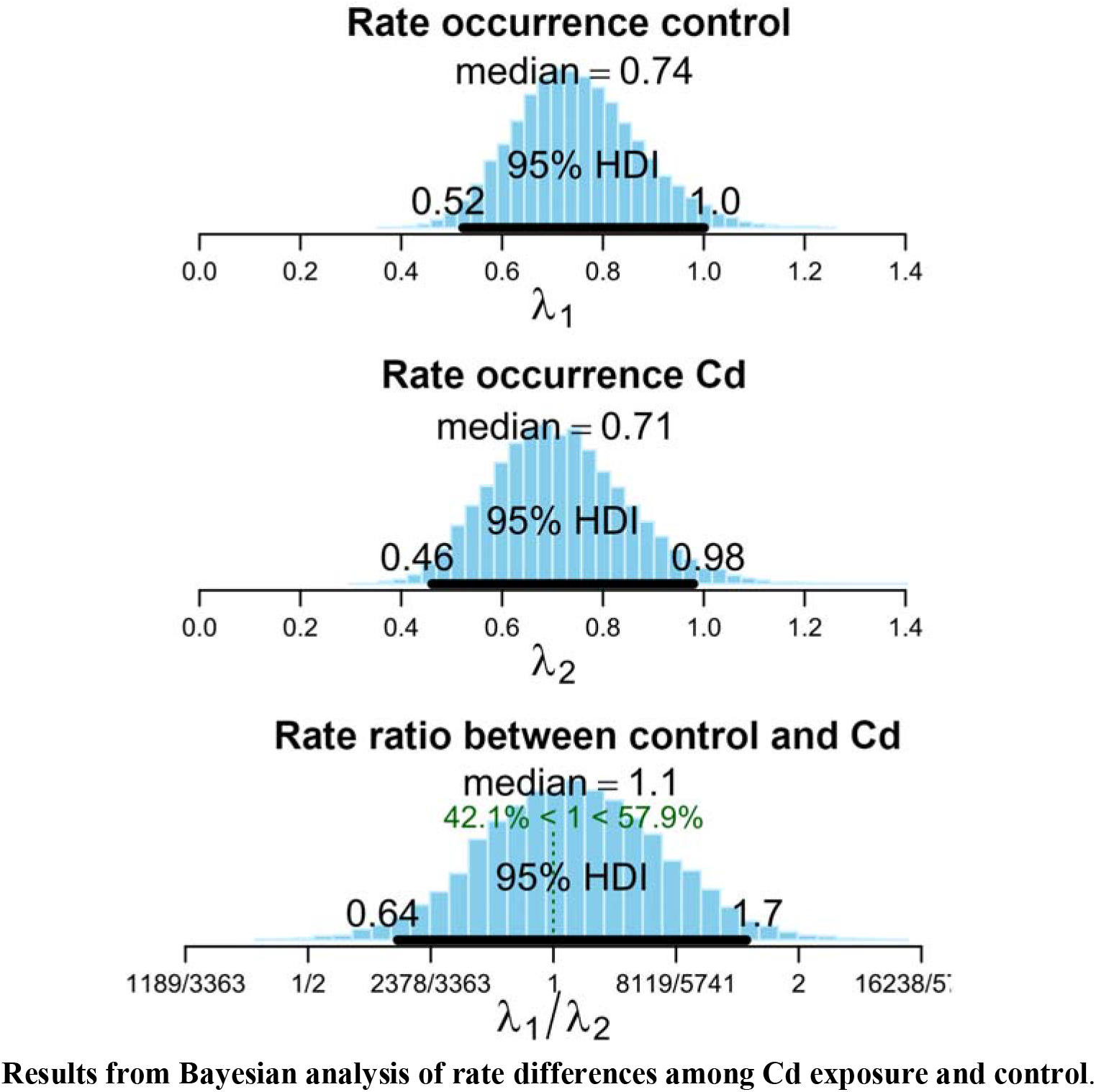
Shown are the posterior distributions for the mutation occurrence per generational passage in control (above) and Cd exposure (middle) and the ratios of the rates (below). The 95% Highest Density Intervals (HDI) are indicated as black bars, respectively. The probabilities for the true rate being larger or smaller than 1 are presented in green in the lower panel. There is no conclusive evidence for either rate being higher than the other.

**Table 1.** Summary information on MA lines. Mean coverage, number of callable sites, number of single nucleotide mutations (SNMs) and resulting mutation rate (μ) per generation for each of the mutation accumulation (MA) lines in *C.riparius.*Results from Bayesian analysis of rate differences among Cd exposure and control.

| Control |  |  |  |  | Cd |  |  |  |  |
| --- | --- | --- | --- | --- | --- | --- | --- | --- | --- |
| MA line | mean coverage | callable sites | SNM | $\mu$ estimation | MA line | mean coverage | callable sites | SNM | $\mu$ estimation |
| 1 | 46.71 | 94074863 | 3 | 6.3779E-09 | 3 | 41.79 | 123248330 | 1 | 1.62274E-09 |
| 2 | 43.61 | 112139472 | 2 | 3.56699E-09 | 6 | 39.77 | 134886417 | 2 | 2.96546E-09 |
| 3 | 37.59 | 144770197 | 3 | 4.1445E-09 | 7 | 38.63 | 138688159 | 6 | 8.65251E-09 |
| 4 | 36.2 | 147108695 | 3 | 4.07862E-09 | 8 | 40.51 | 130790009 | 3 | 4.58751E-09 |
| 6 | 43.59 | 117262208 | 3 | 5.11674E-09 | 10 | 40.16 | 135389419 | 5 | 7.3861E-09 |
| 7 | 41.77 | 120922998 | 5 | 8.26973E-09 | 11 | 36.82 | 145693540 | 3 | 4.11823E-09 |
| 8 | 38.39 | 137175281 | 3 | 4.37397E-09 | 12 | 40.79 | 129861161 | 4 | 6.16043E-09 |
| 9 | 40.3 | 135000368 | 4 | 5.92591E-09 | 13 | 39.85 | 137103976 | 4 | 5.83499E-09 |
| 10 | 40.21 | 136776246 | 4 | 5.84897E-09 |  |  |  |  |  |
| 12 | 40.98 | 128383362 | 7 | 1.09048E-08 |  |  |  |  |  |

### 3.2. Cd influence on mutation spectrum and mutation rate

Cd exposure did not influence the mutational spectrum. In the control group 29 SNMs were transitions, while 8 SNMs were trans vers ions. The Cd exposed group showed a similar trend with 20 transitions and 8 transversions, which was not significantly different (χ^2^ = 0.66667; p= 0.41) (Table 2). The same was true for the ratio of A/T > G/C and G/C > A/T mutations (0.43 in the control versus 0.39 in the Cd treatment, (χ2 = 0.036; p= 0.85).

**Table 2.**
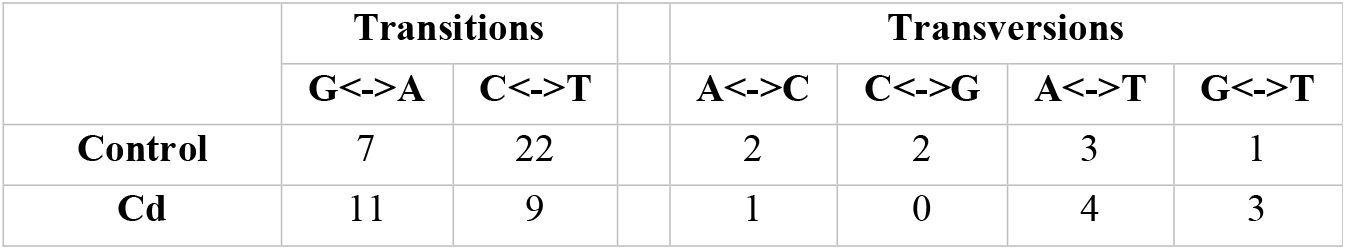
Summary information on mutational spectrum. Number of transitions and transversions and A/T > C/G, C/G > A/T mutations observed on the control and Cd exposed MA lines.

|  | Transitions |  |  | Transversions |  |  |  |
| --- | --- | --- | --- | --- | --- | --- | --- |
|  | G<->A | C<->T |  | A<->C | C<->G | A<->T | G<->T |
| **Control** | 7 | 22 |  | 2 | 2 | 3 | 1 |
| **Cd** | 11 | 9 |  | 1 | 0 | 4 | 3 |

The μ estimate of the haploid SNM rate per generation and site for the control group was μ = 5.86 x 10^−9^ (95% CI: lower = 4.42 x 10^−9^; upper = 7.30 x 10^−9^) and for the Cd group μ = 5.16 x 10^−9^ (95% CI: lower = 3.23 x 10^−9^; upper = 7.10 x 10^−9^, Table 1).

## 4. Discussion

Germline mutagenicity induced by environmental stressors and anthropogenic substances in metazoans is an important issue, but currently largely untested because measuring mutation rates directly was until recently a lengthy, work intensive and costly undertaking. The over-exponential drop of NGS sequencing cost (Shendure and Aiden, 2012), the increase in sequencing accuracy, the introduction of short term mutation accumulation approaches (Oppold and Pfenninger, 2017) and the development of rigorous analysis pipelines (Howell et al., 2018) rendered this venture now reasonably short and economically.

The experimental procedure in producing short-term accumulation lines is mostly straightforward (Foucault et al., 2019). One important issue is the necessity to start with an excess of MA lines (1.5-2 times the intended number), since it is not uncommon that single lines fail to reproduce at some point during the process. Since the lines reproduce with full-sib mating only, it is not unlikely that a deleterious recessive allele present in the parents becomes homozygous in the egg-clutch chosen for lineage propagation.

The most critical issue in the procedure is the automated identification of candidate mutations from the WGS output. Mutations are exceedingly rare (the proverbial needle in a haystack) while WGS data is noisier as one could wish. It is therefore a substantial challenge to reliably and accurately separate the signal from the noise. To the present moment, the majority of MA line studies employed different custom bioinformatic pipelines to identify the *de novo* mutations (Bull et al., 2019; Liu et al., 2017; Oppold and Pfenninger, 2017; Wamucho et al., 2019) making the process of data analysis not only complicate and indirect, but also likely to generate false positives. The use of accuMUlate software, which is flexible to accommodate exclusive properties of the experiments while reflecting the design of a typical MA line design (Howell et al., 2018), makes the data analysis more straightforward, uniform, and consequently, faster. Comparison of the control rate estimated here with the mutation rate independently estimated at the same temperature showed an almost identical rate (unpublished data). Thus, the specifications employed to run the software and the strict filtering (Long et al., 2016), together with individual IGV visualization of the candidates (Keightley et al., 2015), suggested that the pipeline employed for data analysis was robust and accurate enough to yield valid, comparable estimations and the results here presented are likely to be precise. All of this together implies that the present experimental design and its described data analysis are suitable to evaluate the mutagenicity effects of chronic stress generated by environmental pollutants on the evolutionary relevant germline mutation rate over a reasonable period of time.

We used a Bayesian framework to analyse the difference in mutation rates between Cd exposure and control (Bååth, 2014). Bayesian analyses have been repeatedly advocated in ecotoxicology (Billoir et al., 2008; Pollino and Hart, 2005). They are particularly suitable because ecotoxicological assessments are destined to improve decision making, for which quantification of the effect sizes and the uncertainty associated with these predictions are more essential than an arbitrary significance threshold (Feckler et al., 2018). The Bayesian credible intervals of the mutations rate ratios here can be directly interpreted as probabilities of the effect sizes. In the present data (Figure 1), we found no evidence that long-time exposure to environmentally relevant Cd concentration increased the mutation rate.

In the case of Cd, a well characterized environmental pollutant, its use in the manufacturing industry of paints, batteries and plastics and its presence in phosphate fertilizers is causing serious concern regarding its occurrence and effects in the environment (Halwani et al., 2019; Pan et al., 2010). Yet, although Cd is genotoxic, interferes with DNA repair machinery, shows clastogenic effects, provoke DNA strand breakage and exert oxidative stress (Filipič, 2012; Hartwig, 2018; Sherrer et al., 2018), its mutagenesis mechanism is still undefined (Morales et al., 2016). Oxidative stress may contribute to DNA strand breakage that, by its turn, can be misrepaired. Therefore, Cd being a toxicant that acts on either generation of reactive oxygen species and on the DNA repair outcome (Ling et al., 2017; Morales et al., 2016), exposure to it should lead to higher mutation rates (Jin et al., 2003). However, the present research shows that *C. riparius*, chronically exposed to Cd for three consecutive generations, did not show any hints for an increase in mutation rate, nor a change in the mutational spectrum. As discussed by Doria and Pfenninger (2020) the possibility that the organisms were not stressed enough can be discarded, since at the same experimental conditions and Cd concentration, higher mean emergence time and mortality were recorded. But, despite the evident stress at which animals were submitted, *C. riparius* is reported to cope with different Cd exposure scenarios (Gillis and Wood, 2008; Marinkoviċ et al., 2012) and we cannot exclude the need to analyse Cd mutagenicity in other organisms to exclude any species-specific effect. Further, it cannot be excluded that Cd might have mutagenic effects at even higher concentrations, however, as already reported for *Caenorhabditis elegans*, its toxicity may prevent reproduction before mutagenicity sets in (Wamucho et al., 2019). Finally, on natural habitats the organisms will be hardly exposed to just one contaminant, but to a complex mixture of them. Thus, the growing *in vitro* evidence that Cd can act as an mutagenic enhancer ought be investigated (Hartwig, 2018; Mukherjee et al., 2004).

The lack of SNMs rate or spectrum change under stress conditions is consistent with the three previous MA studies involving WGS and multicellular animals (Bull et al., 2019; Joyner-Matos et al., 2011; Wamucho et al., 2019). One reason might be that DNA strand breaks caused by Cd exposure are more prone to be repaired as loss-of-heterozygosity or copy number variants (CNVs) than into SNMs (Lieber, 2011). Moreover, due to Cd tissue-specific affinity and distribution (Swiergosz-Kowalewska, 2001; Hensbergen et al., 2000) in multicellular animals, higher mutation rates might be occurring only in somatic cells. The clear distinction between germline cells that rarely divide and much more divisible somatic cells could lead to a greater susceptibility of the latter (Fitzgerald et al., 2017). Finally, elevated mutation rates may appear only in susceptible genotypes and populations of small effective size in a positive mutational feedback loop (Lynch and Gabriel, 1990; Sharp and Agrawal, 2016).

Consequently, the present study can only supply indirect information regarding Cd environmental exposure on humans. First, humans are more susceptible to Cd via inhalation in occupational exposure scenarios and/or via food, tobacco smoke and ambient air (Filipič, 2012; Hartwig, 2010). Because different routes of exposure are known to, not rarely, exert different genotoxic effects (Vermeulen et al., 2003; Wenberg, 1996) and, second, because of the already proven genotoxicity of Cd, it is not wise to argue against the IARC classification. However, the current study has shown that it may be useful and more precise to clearly distinguish between germline mutagenicity and somatic genotoxicity.

Interestingly, we found two MA lines, whose apparent mutation rate was drastically increased, so-called mutator lines. These two mutator lines occurred only in the Cd exposed group, because of which it might be tempting to ascribe them to the exposure. However, the non-induced, spontaneous occurrence of mutator alleles is known (Raynes and Sniegowski, 2014) and mutator lines have been reported from several direct mutation rate estimations with MA lines (Haag-Liautard et al., 2007) and has been observed on additional studies from our research group in *C. riparius* without exposure to a mutagenic substance. In consideration that the other evaluated replicates did not exhibit the same trend, keeping those MA lines for further analysis would have erroneously altered the estimated mutation rates (Long et al., 2016).

## 5. Conclusion

Chronic multigeneration exposure to low, but environmental relevant Cd concentration did neither affect the basal germline mutation rate nor the mutational spectrum of single nucleotide mutations in *C. riparius.* These not only show that it is impossible to uncritically transfer mutagenicity assessments from unicellular organisms to more complex metazoan animals. Further, we showed that the introduced experimental setup, coupling short-term MA lines with subsequent WGS and a dedicated data analysis pipeline, has the potential to fill the existing gap of evaluation methods for the impact of environmental stressors and anthropogenic substances on mutation rates in multicellular organisms.

## Acknowledgements

The authors thank the LOEWE-Centre TBG funded by the Hessen State Ministry of Higher Education, Research and the Arts (HMWK) for the financial support.

## Declarations of interest

none

## Notes

### Competing Interest Statement

The authors have declared no competing interest.

