## Supplementary material for "Measuring mutagenicity in ecotoxicology: A case study of Cd exposure in *Chironomus riparius*"

S1- AccuMUlate input

nfreqs=0.345 0.155 0.155 0.345

seq-error=0.001

ploidy-ancestor=2

ploidy-descendant=2

mu=2.1e-9

theta=0.001

phi-diploid=0.01

phi-haploid=0.01

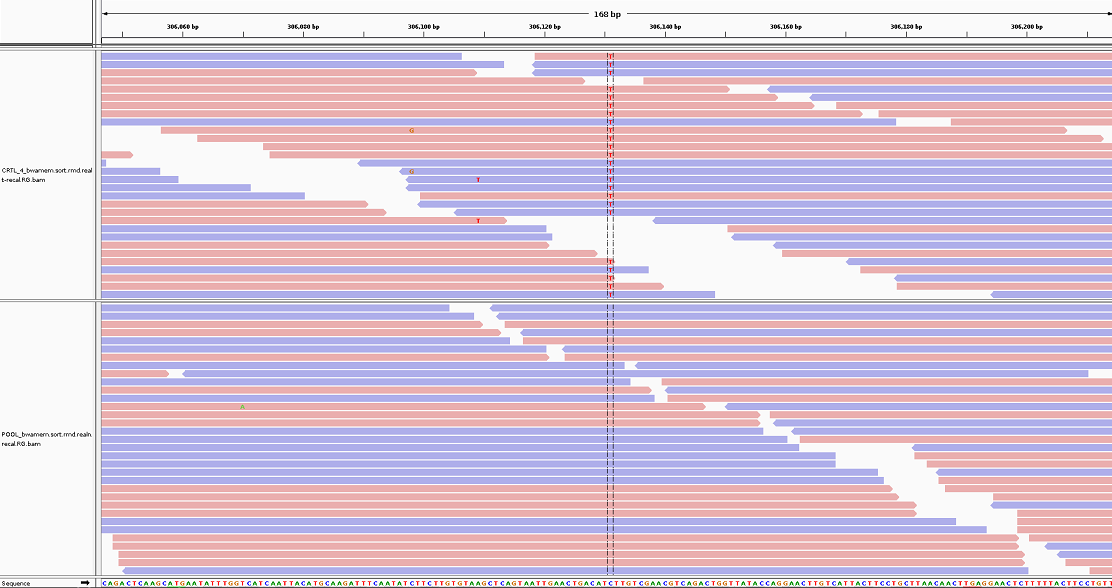
S2 – IGV visualization for final validation

**Figure S2.1** - Screenshot from IGV showing a SNP called (contig: scaffold30_size558987; location: 306131) in individual 4 from the control group (upper panel), which is likely to be a genuine homozygous mutation. Reads from the reference pool (lower panel) are shown for comparison. Pastel red and blue horizontal bars represent sequencing reads for positive rightward (5' to 3') DNA strand and for negative leftward (reverse-complement) DNA strand, respectively. Nucleotides that are different from the reference sequence (shown at the bottom) are indicated.

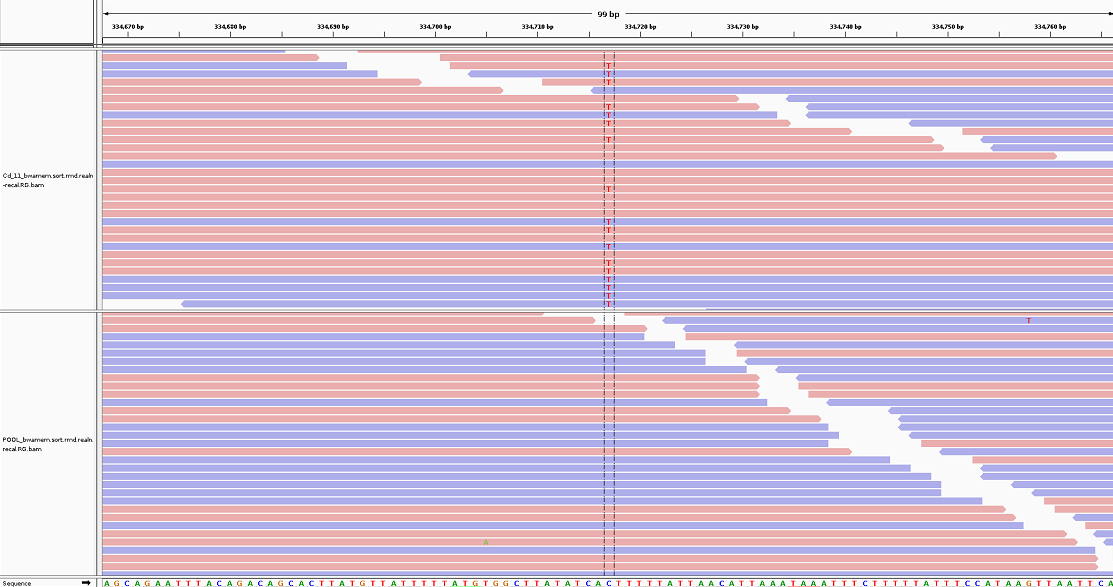
**Figure S2.2 -** Screenshot from IGV showing a SNP called (contig: scaffold81_size1315629; location: 334717) in individual 11 from the Cd exposed group (upper panel), which is likely to be a genuine heterozygous mutation. Reads from the reference pool (lower panel) are shown for comparison. Pastel red and blue horizontal bars represent sequencing reads for positive rightward(5' to 3') DNA strand and for negative leftward (reverse-complement) DNA strand, respectively. Nucleotides that are different from the reference sequence (shown at the bottom) are indicated.

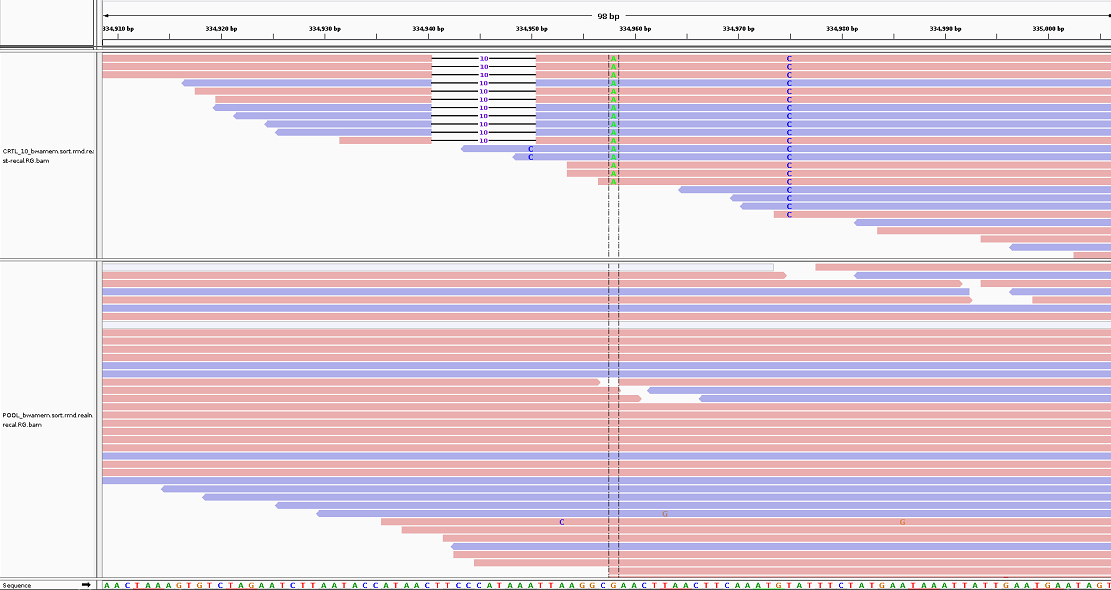
**Figure S2.3** - Screenshot from IGV showing a SNP called (contig: scaffold622_size651815; location: 334958) in individual 10 from control group (upper panel) that is likely to be a false positive, due to deletions and SNPs in association with the candidate mutation not present on the reference pool. Reads from the reference pool (lower panel) are shown for comparison (lower panel). Pastel red and blue horizontal bars represent sequencing reads for positive rightward (5' to 3') DNA strand and for negative leftward (reverse-complement) DNA strand, respectively. Nucleotides that are different from the reference sequence (shown at the bottom) are indicated.

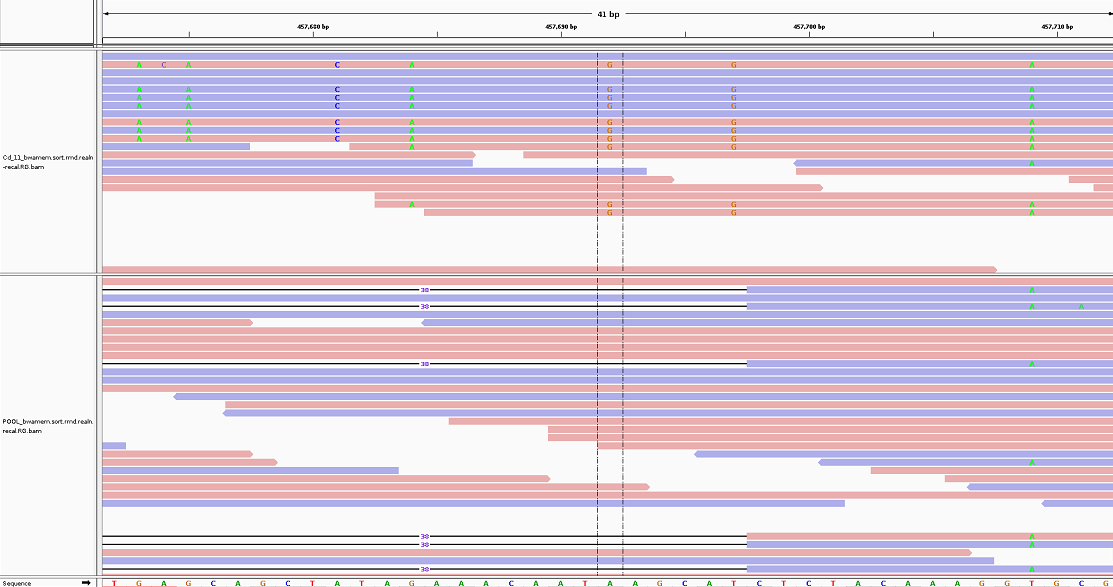
**Figure S2.4 -** Screenshot from IGV showing a SNP called (contig: scaffold249_size746091; location: 457692) in individual 11 from Cd exposed group (upper panel) containing a candidate mutation that is likely to be a false positive, due to the presence of SNPs in complete association
with it that are absent from the reference pool. Reads from the reference pool (lower panel) are shown for comparison (lower panel). Pastel red and blue horizontal bars represent sequencing reads for positive rightward (5' to 3') DNA strand and for negative leftward (reverse-complement) DNA strand, respectively. Nucleotides that are different from the reference sequence (shown at the bottom) are indicated.
